# Rapid antibiotic susceptibility testing and species identification for mixed infections

**DOI:** 10.1101/2021.11.10.468026

**Authors:** Vinodh Kandavalli, Praneeth Karempudi, Jimmy Larsson, Johan Elf

## Abstract

Antimicrobial resistance is an increasing problem globally. Rapid antibiotic susceptibility testing (AST) is urgently needed in the clinic to enable personalized prescription in high-resistance environments and limit the use of broad-spectrum drugs. Previously we have described a 30 min AST method based on imaging of individual bacterial cells. However, current phenotypic AST methods do not include species identification (ID), leaving time-consuming plating or culturing as the only available option when ID is needed to make the sensitivity call. Here we describe a method to perform phenotypic AST at the single-cell level in a microfluidic chip that allows subsequent genotyping by *in situ* FISH. By stratifying the phenotypic AST response on the species of individual cells, it is possible to determine the susceptibility profile for each species in a mixed infection sample in 1.5 h. In this proof-of-principle study, we demonstrate the operation with four antibiotics and a mixed sample with four species.

## Introduction

The rapid increase in antibiotic resistance is a serious threat to human health; access to effective antibiotics is a cornerstone of modern medicine and a prerequisite for e.g. cancer treatment and surgery. Different investigations^1,2^ make different estimations of how grave the situation is but there is a consensus view that action needs to be taken or the costs both in terms of human suffering and global economic impact will be staggering^3^. Experts also agree that the problem is at least partly due to indiscriminate use and misuse of a wide range of antibiotics^4^. To overcome this problem, personalized and rapid antibiotic susceptibility tests (ASTs) are needed, ideally at the point of care^5^. Without these tools, physicians are left with no other option than to prescribe broad-spectrum antibiotics since it can take several days to identify the pathogen(s) and the resistance profile.

The limitations of conventional phenotypic ASTs (disk diffusion agar dilution or broth microdilution) are that they require bacterial growth for extended periods in the presence and absence of antibiotics to see an effect. Using standard culture methods, these tests take one day or more to perform but the time can be reduced to 6-10h with automated systems^6^. However, for certain types of bacterial infections, even a delay of 6h before treatment is initiated can have severe consequences^7^. One such example is sepsis, where the risk of death has been estimated to increase by 7.6 % for every hour that effective treatment is not given^8^. Further, it is shown that in the absence of fast AST, more than 25% of septic patients were treated by clinicians with inappropriate antibiotics, which are strongly associated with mortality^9–11^. Thus, rapid and accurate ASTs are needed to save lives. However, considering the increases in AMR, life-threatening conditions are not the main culprit, but rather the bulk usage of antibiotics for more benign conditions^12^. Resistance is also driven by the strategy to change first-line antibiotics when the local resistance prevalence has reached approx. 10-20%. If fast, reliable ASTs were accessible, high-resistance antibiotics could still be used for the 80-90% of the infections that are still susceptible.

The obvious need and benefits of rapid AST, both for saving lives and guiding prescriptions, have resulted in the development of several new methods over the last decade. These methods are described in a number of recent reviews, e.g. ^4,13^, and we will not repeat all the pros and cons of the different methods here. Briefly, the methods can be divided into genotypic and phenotypic. Genotypic methods identify specific genetic markers that are associated with antibiotic resistance. Although this results in rapid detection of specific resistance genes, it depends on our knowledge of resistance mechanisms which is far from complete^14^,in particular considering the rapid emergence of new resistance mechanisms. Furthermore, the absence of resistance genes does not predict susceptibility to antibiotics ^15^, i.e., you may learn what not to use but not what will work. In phenotypic methods, the bacteria are exposed to an antibiotic and their phenotypic response, e.g. lysis, growth rate reduction, is monitored. Phenotypic methods work irrespective of the mechanism of resistance. If the phenotypic response is there, the bacterium is susceptible. The various rapid phenotypic AST that have reached the market can deliver an answer (susceptible or resistant) in 2-6h for positive blood cultures that have been growing >6h from sampling the patient. For other samples, e.g. urine, the time-to-answer can be reduced to around 30 min by loading the sample directly into a microchip and measuring the growth-rate with and without antibiotics^16^.

A common problem for all rapid phenotypic AST methods that assay the patient sample directly without preculture, is that they only work if the species of the bacteria is known ^17^. For infections with a narrow spectrum of pathogens, this is not a problem, but for sepsis and other more complex infections, species ID is essential. MALDI-TOF mass spec is currently the golden standard for species determination^18^. The technique has until recently only been used for isolated colonies on primary plates, but the practice is changing for blood cultures to remove the additional time of culturing on plates (12h) after the blood culture^19^. MALDI-TOF does however require pre-culture of single bacterial species and may not work well for mixed infections commonly seen in sepsis, wounds, catheter-associated UTIs^20^, and community-acquired pneumonia among others^21^. The challenge remains to make rapid phenotypic AST with species ID that can be used directly on patient samples without a preculture step.

To address this issue, we use a microfluidic chip that is capable of rapid capture of individual bacteria from a sample and allows optical monitoring of their growth with and without antibiotics. Next, we identify the bacterial species by fluorescence in situ hybridization (FISH) with species-specific ssDNA probes targeting the 16s/23s rRNA sequence. Once we have the species ID and AST response for each position in the microfluidic chip, we stratify the AST response based on species. A schematic overview of the approach is presented in figure 1.

**Figure 1:**
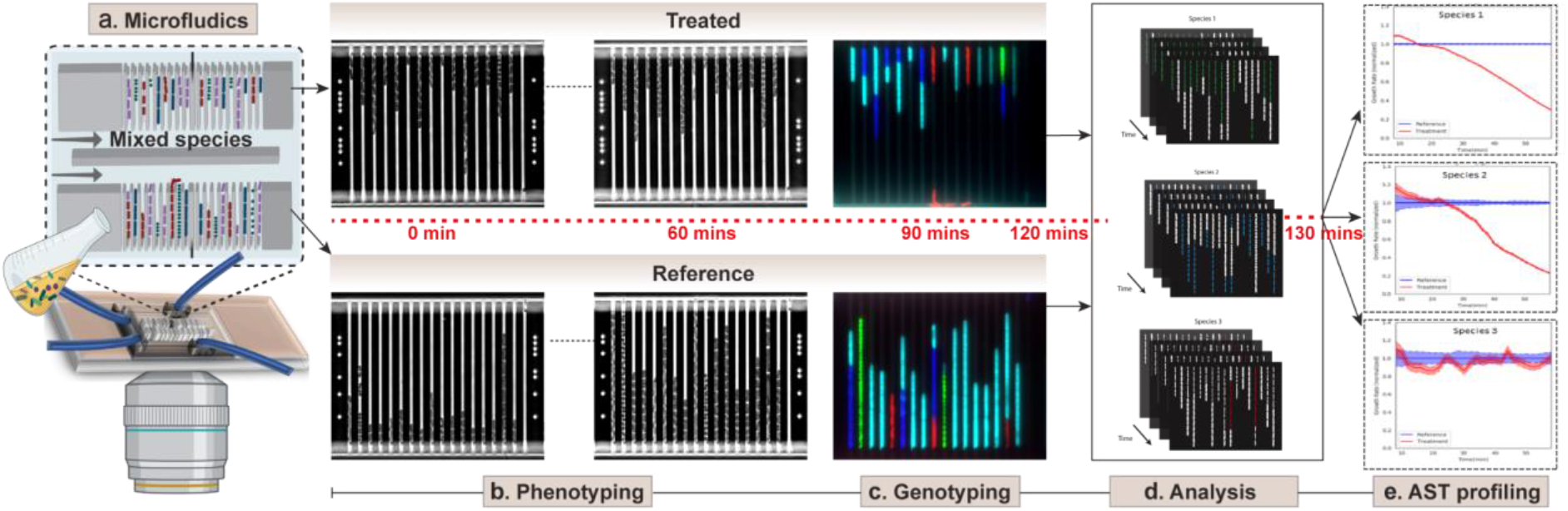
Schematic representation of the AST workflow with timeline. a) A cartoon of the microfluidics setup with the mixed species loaded on the chip. b) Time-lapse phase contrast images of the cells in the traps when grown in media with (top) and without (bottom) antibiotics. c) Fluorescence images of the bacteria with ssDNA probes targeting the ribosomal RNA of specific bacteria for species identification. d) Analysis of time-lapse stacks and species ID using deep learning for segmenting and tracking cells. e) Detection of AST profiles for individual pathogens at a given antibiotic concentration.

In this proof of principle application, we perform the ASTs for four common pathogens (*Escherichia coli*, *Klebsiella pneumonia, Pseudomonas aeruginosa* as examples of gram-negative strains and *Enterococcus faecalis* as an example of a gram-positive strain) and four different antibiotics from different classes: Vancomycin (Van) [Glycopeptide], Ciprofloxacin (CIP) [Fluoroquinolones], Gentamicin (Gen) [an aminoglycoside], and Nitrofurantoin (NIT) [other agents]. In the end, we show how single-round, multi-color labeling enables identification of up to ten species simultaneously.

## Results

### Phenotypic AST followed by Genotyping by FISH

The culture chip that we have developed for this assay is capable of rapid capture of bacteria directly from a liquid sample and allows for optical monitoring of the bacterial growth with and without antibiotics in real time. The same chip design has previously been used to capture bacteria from blood cultures despite an overwhelming excess of blood cells^22^. The chip features two rows of 3,000 cell traps each. Each trap measures 1.25 × 1.25 × 50 μm^16^ with a constriction at the end which prevents the bacteria from escaping the trap, while still allowing media and probes to flow around the cells. To simulate a mixed infection situation, bacterial overnight cultures of four different species were diluted in a Mueller Hinton (MH) broth, pooled, and directly loaded in the microfluidic chip. In a typical experiment, loading one or a few cells in each trap takes 1 min at ∼10^5^ CFU/ml. We supplied growth media with antibiotics to the traps in one of the two rows and plain growth media to the other.

The phenotypic response to the antibiotic was determined in ≈60 minutes by capturing the phase-contrast images of each cell every 2 min and calculating the growth rates of individual cells. The phenotypic response can be pushed to shorter times depending on which antibiotics are used and at which concentration^16^. To identify the species of each bacterial cell, we performed fluorescence *in situ* hybridization (FISH) using species-specific, fluorescent ssDNA probes. These probes bind to the very abundant 16s/23s ribosomal RNA sequences (Supplementary Table 1) and have previously been successfully used for species identification in positive blood cultures^23^. The species classification method for FISH signals is described in SI section 3.

### Analysis using Deep learning models

Performing growth rates analysis on multi-species samples requires a general method for detection and tracking of cells that come in different shapes and sizes. It has recently been shown that U-net architecture^24^ can be used to segment and track *E*.*coli* cells^25^ on phase-contrast images with very high accuracy. We performed cell segmentation using the same architecture, trained on *E*. *coli* and *P*. *aeruginosa* phase-contrast images paired with ground-truth segmentation masks determined with fluorescent cells (Fig. 2a). In addition to segmentation masks, weight maps (SI fig 1a) were generated to increase the accuracy of cell boundaries between adjacent cells. The training data was heavily augmented to force the network to learn the invariant features that describe a cell on a phase-contrast image. The network performed well on previously unseen species with different shapes and sizes (SI fig. 2). An overlay of a predicted segmentation mask on a phase-contrast image with mixed species is shown in figure 2b.

**Figure 2:**
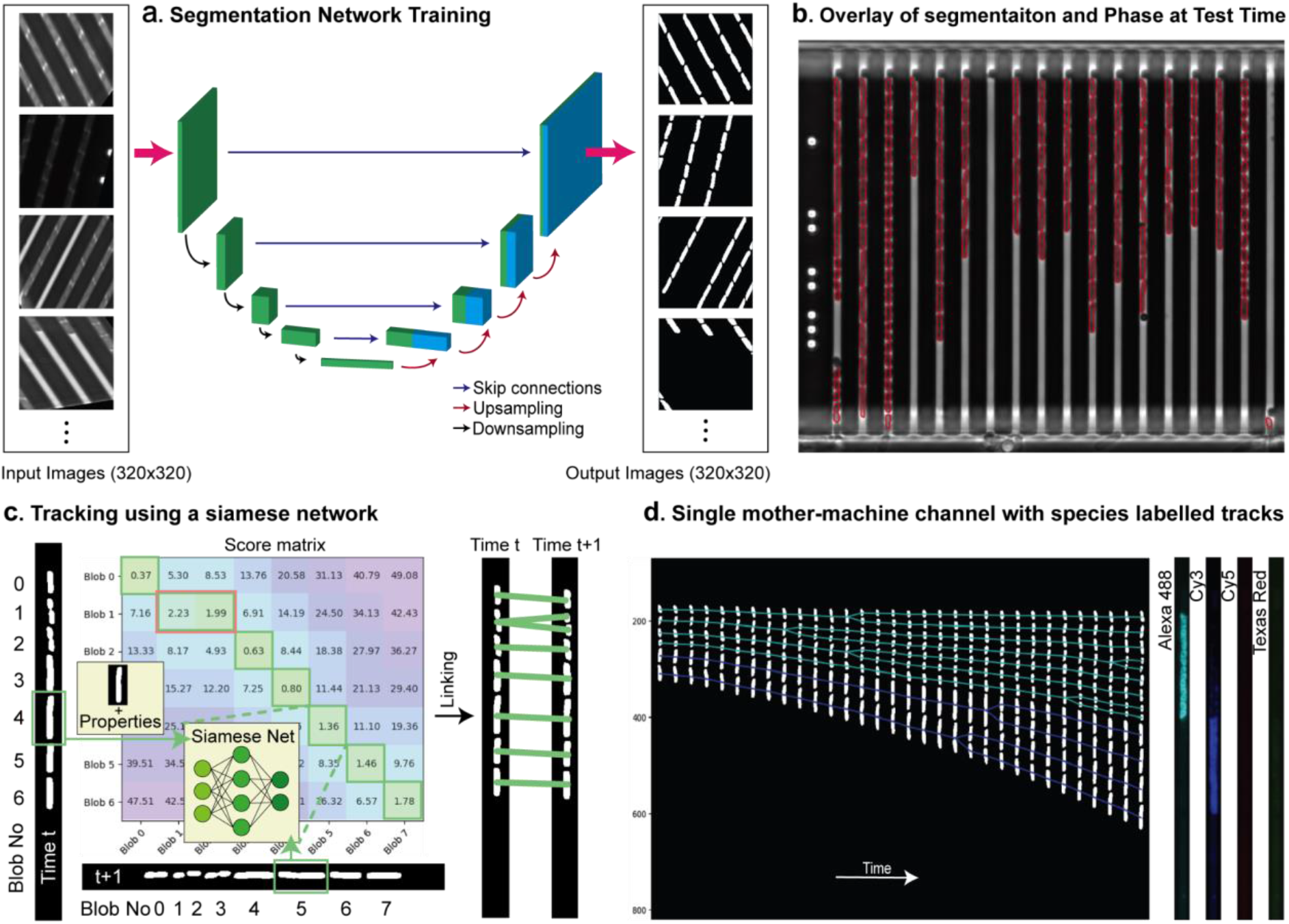
Analysis a) U-net architecture trained for cell segmentation with corresponding input-output pairs of images. b) Overlay of contours from the segmentation mask on a phase-contrast image containing multiple species. c) Each blob from an image at time point t is compared against all blobs in the next frame at time t + 1 to obtain a similarity score (left). Links between two consecutive frames based on the similarity scores (right). d) A timelapse stack of segmented cells in a single mother-machine channel and its corresponding fluorescence images in all four imaging channels. Regions of signal in each fluorescence image are highlighted with a bounding box (red). Tracks in the last frame falling inside these bounding boxes are labelled with corresponding species and propagated backward to time t = 0. E. faecalis tracks with FISH probe in Alexa 488 channel are labeled in cyan and E.coli tracks with FISH probe in Cy3 channel are labeled in blue.

Previous approaches to successful *E*. *coli* tracking have constructed cell-lineages using scoring mechanisms based on overlapping regions^19^ or mother-daughter binary mask predictions using full features from the images^25^. The former approach is very sensitive to tuning overlap parameters while the latter is computationally expensive when a lot of small cells are present in the data. We developed a hybrid approach that performs cell-tracking using a Siamese network^26^ that scores the similarity between cells from one frame to the next (Fig. 2c). The training of both the segmentation network and the tracking network is described in the supplementary information (sections 1 and 2). Species assignment to tracks was performed after cell-tracking (SI section 5). Figure 2d shows a single segmented mother machine trap tracked through time and the corresponding species-labeled tracks.

### Species-wise AST response

With the species ID and AST response for each position in the microfluidic chip, the species-specific AST response could be determined in the mixed samples. Here, we demonstrate the capability of the method to characterize a mixed sample of four different species, although clinical patient samples are more likely to contain only one or two ^27–29^. The AST responses are shown in figure 3a-d. In each experiment, we obtained growth-response curves for three gram-negative strains (*Escherichia coli*, *Klebsiella pneumoniae*, and *Pseudomonas aeruginosa)* and one gram-positive strain (*Enterococcus faecalis)*. The experiments were performed with four different antibiotics: Vancomycin (Van) [Glycopeptide], Ciprofloxacin (CIP) [Fluoroquinolones], Gentamicin (Gen) [an aminoglycoside], and Nitrofurantoin (NIT) [other agents]. We present the results as response plots from individual experiments to simulate the clinical sample situation and, for comparison, we also display the average responses that would have been the result of the growth-rate measurements without species information. Successful AST-profiling could be achieved with samples containing as little as 100 bacteria/species. We used bacteria without specific resistance genes, but since some species have a natural resistance to specific antibiotics, the AST response varies with the species. For example, *Pseudomonas* species are naturally resistant to Ciprofloxacin (Fig. 3a) and Nitrofurantoin (Fig. 3b), and their growth remained unaffected in the presence of 1ug/ml Ciprofloxacin or 32ug/ml Nitrofurantoin. The growth rate of the other bacterial species, on the other hand, dropped more than 10% in 30 min in the presence of these drugs. We also see from the average, non-species-stratified data that without access to species ID, we would not have been able to detect the resistant *Pseudomonas* in the mixed population. Similarly, as expected, all species but *E*. *faecalis* were found to be susceptible to Gentamicin (2μg/ml) (Fig. 3c) and Vancomycin (4μg/ml) (Fig. 3d).

**Figure 3:**
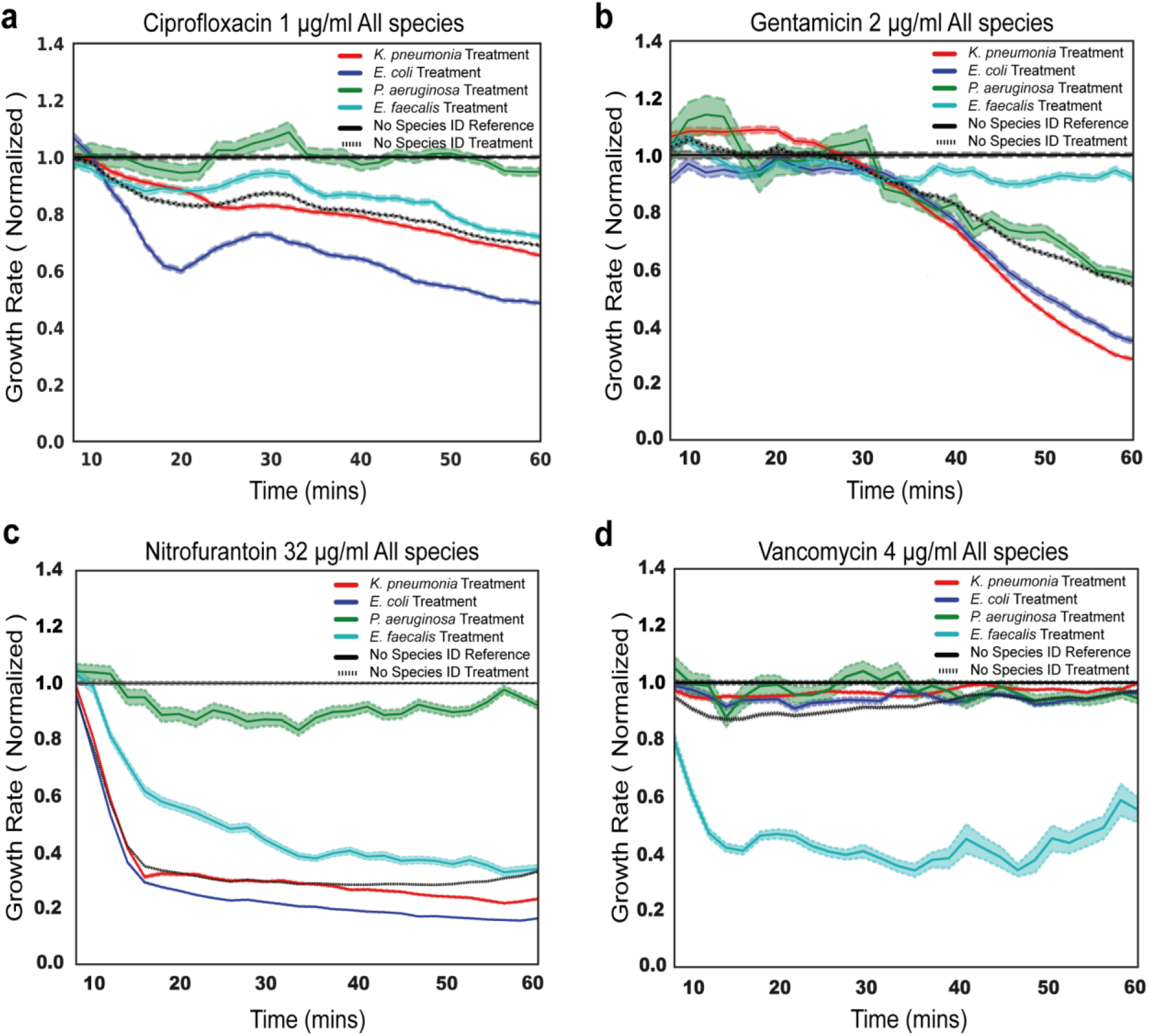
Species-wise response to antibiotic treatment. (a-d) AST profiles with normalized growth rates for the four antibiotics used. The species stratified responses (average and SEM) as well as the pooled response (without species stratification) is shown for each antibiotic.

### Scaling FISH probing up to 10 species

It is estimated that >90% of clinical sepsis samples feature a subset of the 10 most frequent bacterial pathogens^30^. To increase the number of species that we can identify, we performed combinatorial FISH using a species-specific adaptor that can bind two different fluorescent oligo probes. With probes of four different colors, this set-up can identify up to 10 species (Fig. 4a). We demonstrated the combinatorial FISH approach to identify four species loaded in the culture chip (Fig. 4b). Assignment of species was done using k-means clustering on the intensity signal from all four fluorescence channels. Species labels were assigned to clusters manually based on combinations of signals (SI fig 7). Figure 4c shows projections of the clusters in space spanned by the first two principal components of the intensities. Supplementary information section 6 describes this species assignment in detail.

**Figure 4:**
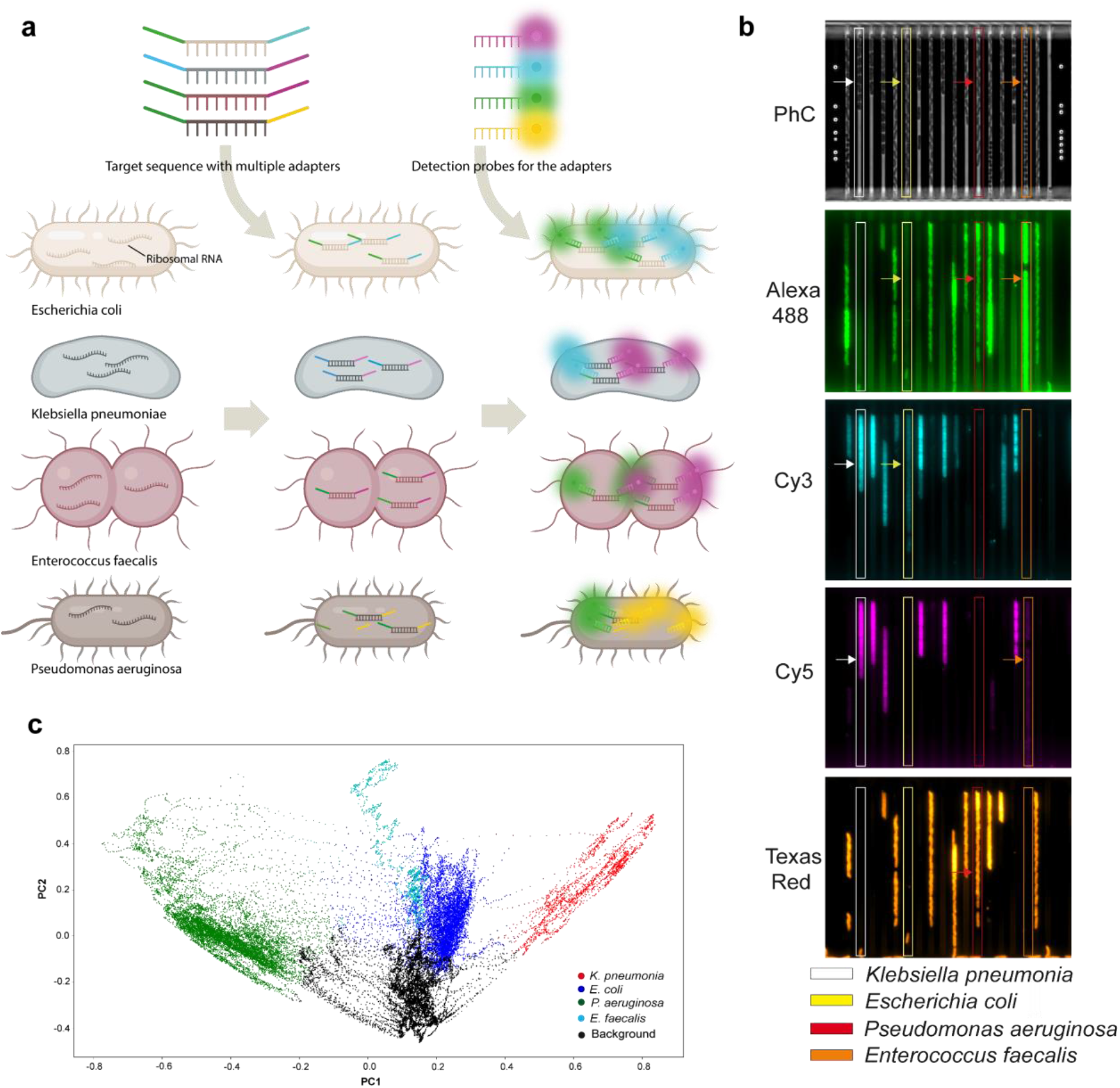
Combinatorial FISH: a) Overview of the combinatorial FISH probing for the multi species identification. A cartoon illustrating the different bacterial species with their ribosomal RNA (left). Illustration of the specific sequences with the multiple adapters targeting the ribosomal RNA of individual bacteria and its hybridization to the target rRNA (middle). Detection probes with different fluorophores. Hybridization of detection probes to the adapter sequences along with unique sequences that are targeted to the species specific rRNA (Right). b) Example images of mixed species loaded in the microfluidic chip and probed using combinatorial FISH for species identification. After the hybridization step, cells were imaged in different channels (PhC, Alexa 488, Cy3, Cy5 and Texas Red). The bacterial species are marked in yellow (Escherichia coli), orange (Enterococcus faecalis), red (Pseudomonas aeruginosa) and white (Klebsiella pneumoniae). c) Clustering of fluorescence imaging intensities using k-means and PCA for species assignment.

## Discussion

In conclusion, we have demonstrated that it is possible to make rapid AST for mixed species infections by performing sequential single-cell phenotypic susceptibility testing and fluorescence *in situ* hybridization in a microfluidic chip. Importantly, ID determination is also applicable for non-mixed infections where it is important to know which MIC-breakpoints to use when making the sensitive-intermediate-resistant call. Although species determination by MALDI-TOF is possible for non-mixed infections and far superior regarding species coverage, this method is limited to centralized hospital labs and samples that are cultured to sufficient biomass. The exact number of cells needed for a MALDI-TOF analysis is hard to find, but it will be orders of magnitude more than what is needed for AST by direct imaging of bacteria. ID with MALDI-TOF will also never be available at the point of care where the majority of antibiotics are prescribed. To guide the initial prescription, it is critical to have the rapid AST as well as sufficiently accurate presumptive species identification near the patient.

Since antibiotic susceptibility breakpoints of closely related species are often the same, it can sometimes be enough to distinguish the bacterial family or genera. For example, an Enterobacteriales-specific sequence can cover *Escherichia, Klebsiella*, and *Salmonella*, which have very similar resistance breakpoints^31^ . This generalization reduces the number of specific probes needed to cover all potential species that can be present in a particular infection. However, if more than 10 specific IDs are required, stripping and reprobing allow for an exponential increase in how many classes can be identified^32,33^, but at the expense of time.

In the current implementation, we ran the test one concentration at a time on a high-end research microscope. To make the technology useful in a clinical setting, the fluidic chip should be parallelized to run multiple antibiotic concentrations simultaneously and with a higher level of automation. Although a minimum of five different concentrations is needed to decide a minimal inhibitory concentration, a simple susceptibility or resistance call can be based on a measurement at a single concentration if the response is well-calibrated to many clinical isolates.

## Methods

### Bacterial strains and Antibiotics

In this study, as a gram-negative representative we used *E*. *coli* K12 MG1655, *K*. *pneumoniae*, and *P*. *aeruginosa*. As a gram-positive representative, we used *E*. *faecalis*. We also used the fluorescently tagged *P*. *aeruginosa* cells, a kind gift from Oana Ciofu^34^. Antibiotics (Ciprofloxacin, Gentamicin, Nitrofurantoin, and Vancomycin) were purchased from Sigma-Aldrich. Stock solutions were prepared as per the supplier guidelines and stored at −20°C. The solutions were thawed to room temperature before performing the AST experiments.

### Media and culture conditions

In all experiments, Mueller-Hinton (MH) medium (70192; Sigma-Aldrich) was used as a broth. Overnight cultures (ONC) were prepared by inoculating bacteria from glycerol stocks (−80 °C) in MH medium and incubating at 37 °C for 14-15 hours with continuous shaking (200 rpm). From ONC, cells were diluted 1:1000 times into fresh MH medium supplemented with a surfactant (Pluronic F-108; 542342; Sigma-Aldrich; 0.085% (wt/vol) final concentration). The liquid culture was grown by shaking at 200 rpm at 37 °C for 2 hours. Next, to perform AST experiments, we mixed the different strains in equal concentrations and loaded them on the microfluidic chip.

### The Microfluidic chip and setup

The chip consist of mainly two parts: a micromolded silicone elastomer [Sylgard 184; polydimethylsiloxane (PDMS)] and a 1.5 mm glass coverslip (Menzel-Gläser) which are covalently bonded together. Chip design and preparation was previously described in^16,35^. Reference^16,35^ describes the numbering of the ports used below. After punching the ports on the chip, it was placed on the microscope and tubing (TYGON) was connected with a metal tubing connector. Briefly, cells were loaded using the port 8.0 and port 2.0 was used for the exchange of medium with the probes. Ports 5.1, 5.2, and 6.0 were used for maintenance of back channel pressure of 500 mbar, ports 2.1 and 2.2 were used for the supply of MH medium with and without the antibiotics at a pressure of 200 mbar. The pressure was controlled by an OB1-Mk3 pump from Elveflow.

### Microfluidic experiments

Imaging starts within five mins after the supply of medium with and without the antibiotics to the cells. We used a Nikon Ti2-E inverted microscope equipped with a Plan Apo Lambda 100x oil immersion objective (Nikon). Images were captured by the Imaging Source (DMK 38UX304) camera. For phase contrast and fluorescence images, we used the optical setup as previously described in^16,35^ and controlled it by Micro-Manager^36^, the in-house build plugin. We maintained 30°C as an optimal condition for cells using a temperature controllable unit and a lexan enclosure (Oklab).

### Fast phenotypic AST

Once the cells were loaded on the chip and exposed to growth media with and without the antibiotics, a total of 80-90 positions on each row on the chip were captured in the phase contrast channel (30 ms exposure time) every two mins for an hour.

### Genotyping

After phenotyping, the medium from the ports 2.1 and 2.2 were depressurised to zero. To fix the cells, formaldehyde (4%) was added by opening, switching the medium and applying pressure 200 mbar from port 2.0 for 10 mins and subsequently washing the cells with 1 × phosphate buffered saline (PBS) for 5 mins. To permeabilize, cells were treated with 70% EtOH for 10 mins and washed with 1 × PBS (5 mins). Permeabilization of gram-positive cells was done by adding the lysozyme (2 mg/ml) for 3 mins and followed by quick washing with 1 × PBS for another 5 mins. For species identification we pooled all specific ssDNA probes (0.1μM) (supplementary table 1) in a hybridization solution (30 % formamide and 2 × SSC) and hybridized for 30 mins at 30°C. Next, we captured the fluorescence images for each individual probe in different channels (TYE 665, TYE 563, Texas Red and Alexa Fluor 488) at 300 ms exposure times and respective phase contrast images at 30 ms exposure time. In total, it took 20 mins to image all the positions in all the channels on the chip.

### Cell segmentation and Channel Detection

Phase-Contrast images of the cells growing in channels were segmented for both cells and channels using a deep learning model with U-net architecture. The cell-segmentation model was trained with data obtained from imaging *E*. *coli* (K12 MG1655 intC::P70-venusFast) and *P*. *aeruginosa* strains (PAO1-mCherry-P_cd_-GFP+) that constitutively express mVenus and GFP respectively. The training procedure was enhanced with data augmentations to force learning features to discriminate cells from the background. The model training and performance on unseen species are described in SI Section 1. The channel segmentation model was trained with data that was refined based on histogram profiles, also described in SI Section 2. After obtaining channel locations, time-series stacks of segmented cells and corresponding fluorescent channel images were bundled for tracking, species assignment, and growth rate calculations.

### Cell Tracking

Cells were tracked through time using a neural network that scores similarities between cells from one frame to the next. This neural network has a Siamese net architecture and was trained on a few time-series stacks of multispecies cells that were manually annotated (SI section 4). Cells between frames were linked based on the scores and tracks were generated by chaining a series of links. Cell tracks were corrected for errors (SI section 4).

### Species assignment and Growth Curve splitting

The fluorescent images for each mother machine channel were corrected for background and continuous regions of signal above thresholds were mapped to species labels (SI section 3 and 5). Regions with multiple fluorescent labels for one species were classified using k-means clustering and PCA (SI section 6). Cell tracks falling in these regions in the last frame before fixing cells were labeled with the corresponding species and all the species labels were rolled back to time point 0. Growth rates were calculated by fitting exponential curves on the areas of cells in a moving 5 timepoint window (SI section 7).

### Oligos and Probes design

FISH probe sequences for the individual target rRNA were obtained from the Probase^23,37^ and purchased from the Integrated DNA Technologies (www.idt.com), see supplementary table 1. For the combinatorial FISH probing, we used barcode sequence and detection probes, which are listed in supplementary table 2-4, purchased from IDT.

## Acknowledgements

We wish to thank I. Barkefors for helpful input on the manuscript and help with the figures. We acknowledge funding from SSF (ARC19-0016), the European Research Council (BIGGER:885360), the Knut and Alice Wallenberg Foundation (2016.0077; 2017.0291; 2019.0439) and the eSSENCE e-science initiative to J.E.

## Author contributions

J.E. conceived the *in-situ* ID after AST method, P.K. developed and implemented the analysis methods and analysed the data. V.K and J.L performed the experiments and developed the protocol for FISH probing in the fluidic chip. J.E., P.K. and V.K. wrote the paper.

## Materials & Correspondence

All strains, raw data and analysis codes will be provided upon reasonable request to J. E.

