## Supplementary material for "Rapid antibiotic susceptibility testing and species identification for mixed infections": Supplemantary methods

##### Section 1: Cell Segmentation: U-net training and performance

The cell segmentation training data contains 3056 phase-contrast and the corresponding fluorescence images of *E.coli* expressing the fluorescent marker mVenus (2583 images) and *P.*aeruginosa (473 images) expressing the fluorescent marker GFP. Binary masks were obtained by denoising and performing adaptive (otsu) thresholding on the fluorescent channel images. The width and height of these images are 1042x856 pixels and a total of 2445 images were used for training. The U-net was trained on images of size 320x320 pixels. The number of training examples used during one iteration of the training loop, i.e. batch size is 8. Each of these images were obtained by randomly cropping 320x320 pixels from the original images. These cropped images were scaled with a random ratio between 0.75 and 1.25, to cover various ranges of cell sizes and resized back to 320x320. Random noise, illumination adjustments, gaussian blurring, and rotation of the images were done on the training data pairs as described in[1]. Any operation that gave a non-binary value in the binary mask was thresholded using otsu thresholding. 20% percent of the training data was used for validation of the model. In addition to image augmentations, weight maps were generated using morphological operations to give more weight to the pixels separating cells in the loss function, as described in the original U-net paper[2]. **SI fig 1a** shows a sample input image, its corresponding binary mask and weight map used during the training process. The loss function used to train the net is dice-loss + weighted binary cross-entropy[3].

Adam optimizer [4] was used with a learning rate of $10^{-4}$ and the U-net was trained for 10 epochs until the losses flattened out as seen in **SI fig 1b**. The model learnt features used for discriminating cells from the background. This can be seen from segmentation masks generated on previously unseen species E. *faecalis* (**SI fig 2c**). The model output is a probability map of each pixel being a cell or not. This output was thresholded to obtain a binary mask of the segmented cells. During test-time, raw phase-contrast images of size 2048x1024 pixels were fed directly to the net to obtain cell segmentation masks. The full phase-contrast image and the corresponding output from the network after thresholding is shown in **SI fig 1c**.

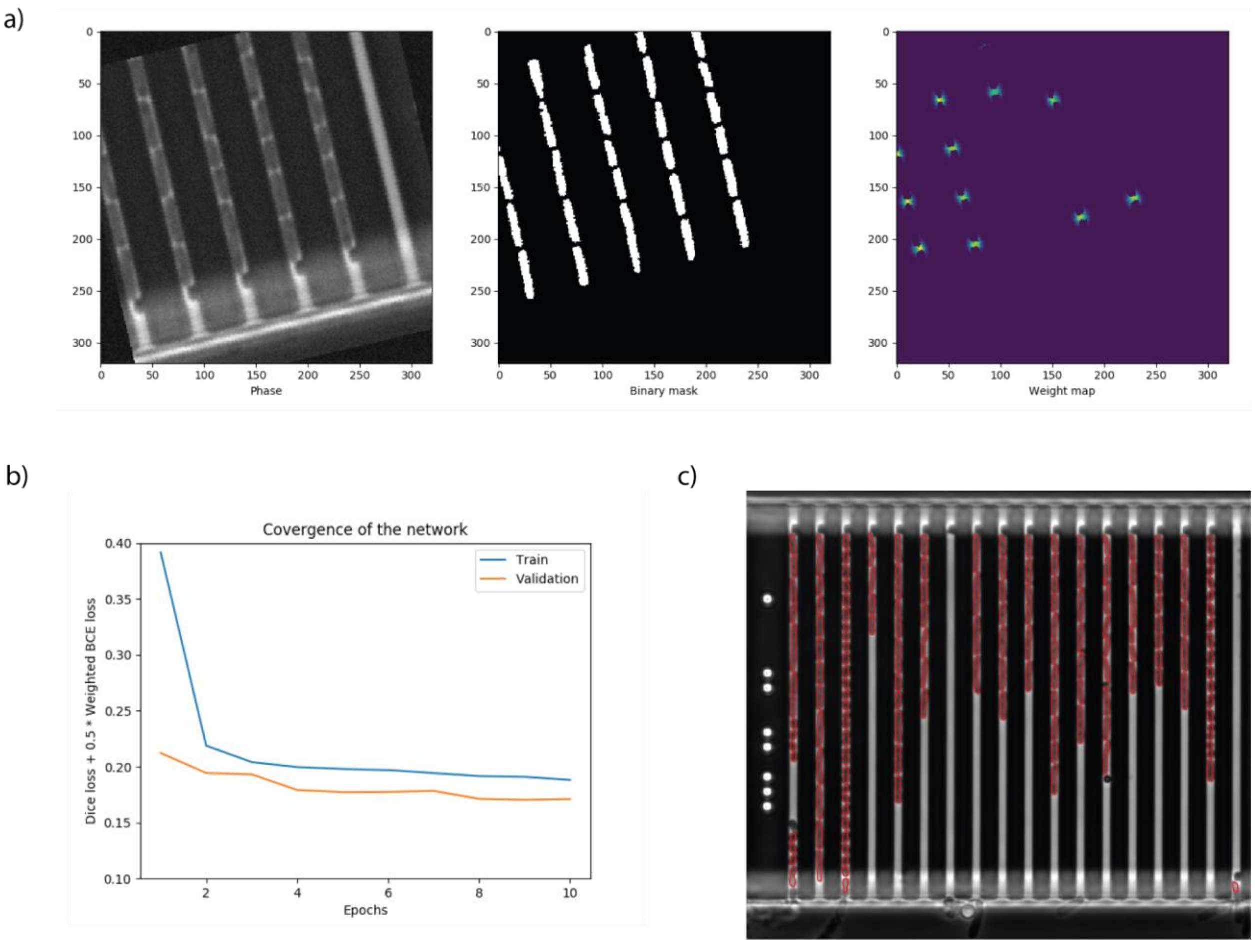


***Supplementary Figure 1:*** *a) Cropped phase-contrast image (left) and it's corresponding binary mask from thresholded fluorescence image(middle) and weight map(right) b) Training and validation loss curves. c) Phase-contrast image with overlay of contours from the cell segmentation.*

**SI figure 2** shows the overlay of segmentation mask on phase-contrast images for each individual species loaded in separate chips. The same network was used on all images and the data has not been part of training of the network.

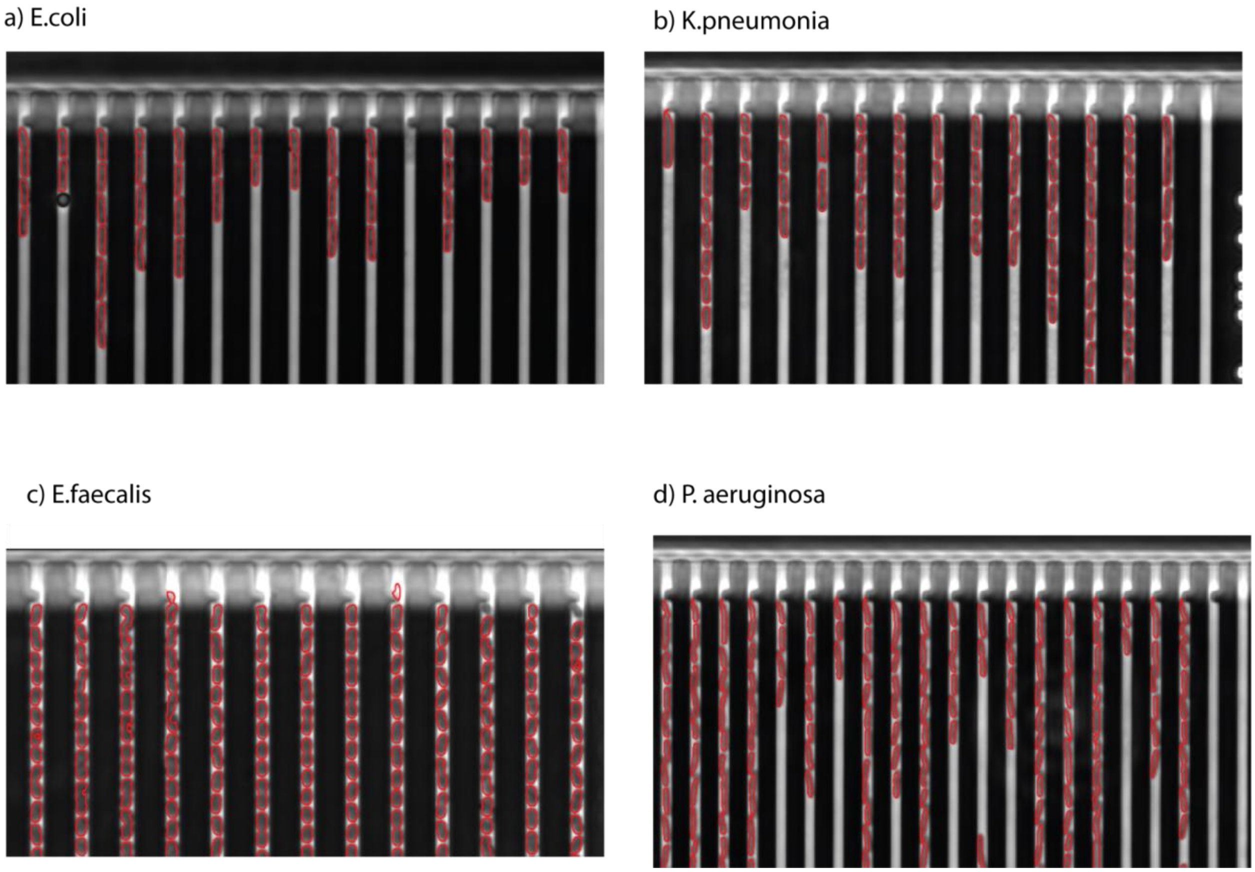


***Supplementary Figure 2:*** *Segmentation mask from the cell segmentation network overlayed on phase-contrast images of individual species images. A) E. coli B) K. pneumoniae C) E. faecalis D) P. aeruginosa*

### Section 2: Channel detection

Channel detection was done using the same U-net architecture as used for cell segmentation. The original U-net described in[2] has the number of feature channels varying from 64 at the input to 1024 before up sampling, doubling after every down sampling step. In this channel detection net, the number of the feature channels were reduced by a factor of 8 at all stages compared to the original U-net model. Training data was generated using image processing operations (histograms of intensity, peak finding on intensity histograms, binary morphological operations, etc.). The same data augmentations and loss functions used for the cell segmentation model were used during training of the channel detection network (**SI section 1**). All pixels were given the same weight in the loss function. The training optimization is similar to that of the cell segmentation model training. **SI fig 3a** shows a sample of the training data used as input for this net. **SI fig 3b**, shows the training and validation loss curves and **SI fig 3c** shows the raw phase-contrast image overlayed with contours of the channel segmentation.

The mother machine channels were detected in each phase contrast image of the time-lapse microscopy data and time-series stack of each individual channel were constructed for further analysis.

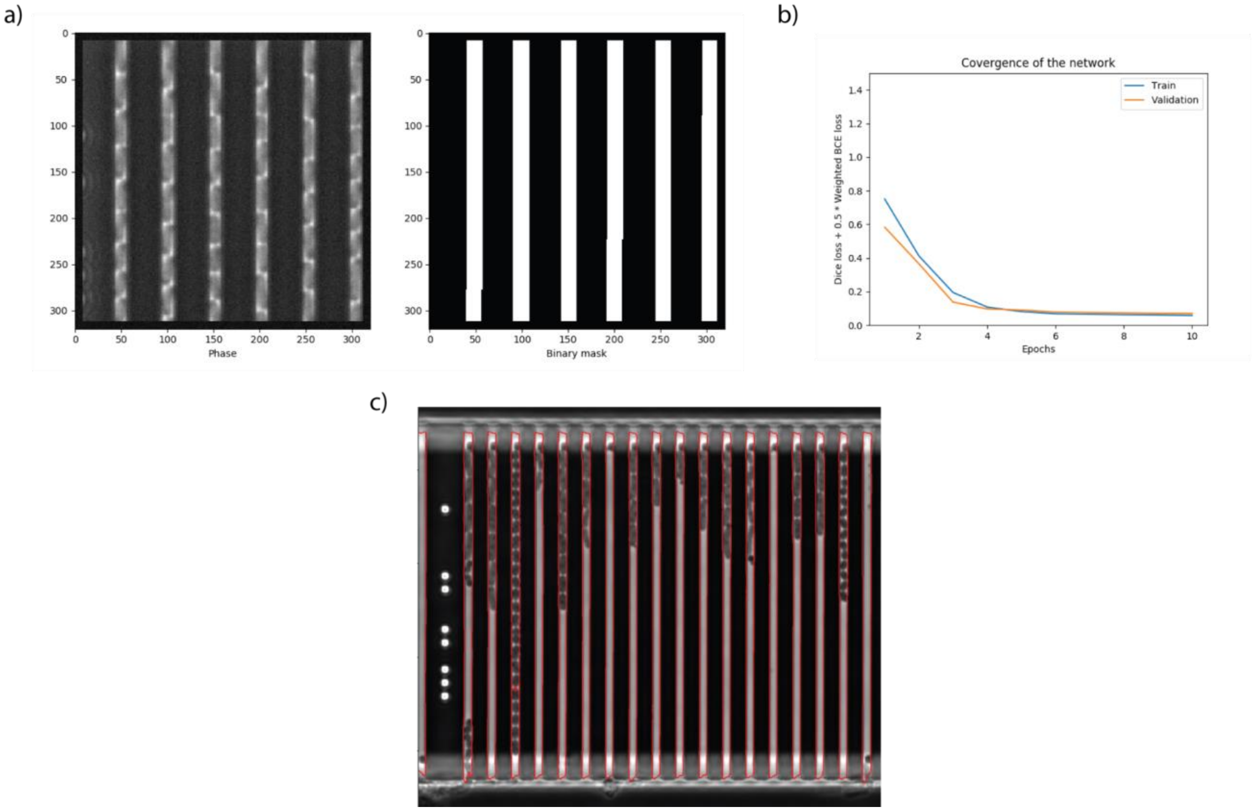


***Supplementary Figure 3:*** *a) Sample training data b) Training/validation loss curves c) Phase-contrast image and it's corresponding channel segmentation mask*

### Section 3: Processing multi-color FISH Images

Fluorescence images were acquired in 4 channels for the FISH-ID. Each FISH probe (**SI Table 1**) was designed to bind to a single species. Image processing operations like smoothing, histogram equalization was performed on the images and the background was subtracted using the empty channel values present in each image. Regions on each image containing signals above a threshold were detected and regions longer than 50 pixels were considered for species ID. The thresholds were set for each of the 4 fluorescence channel images individually during analysis. **SI fig 4** shows all the 4 fluorescent channel images with bounding boxes outlining the regions with signal.

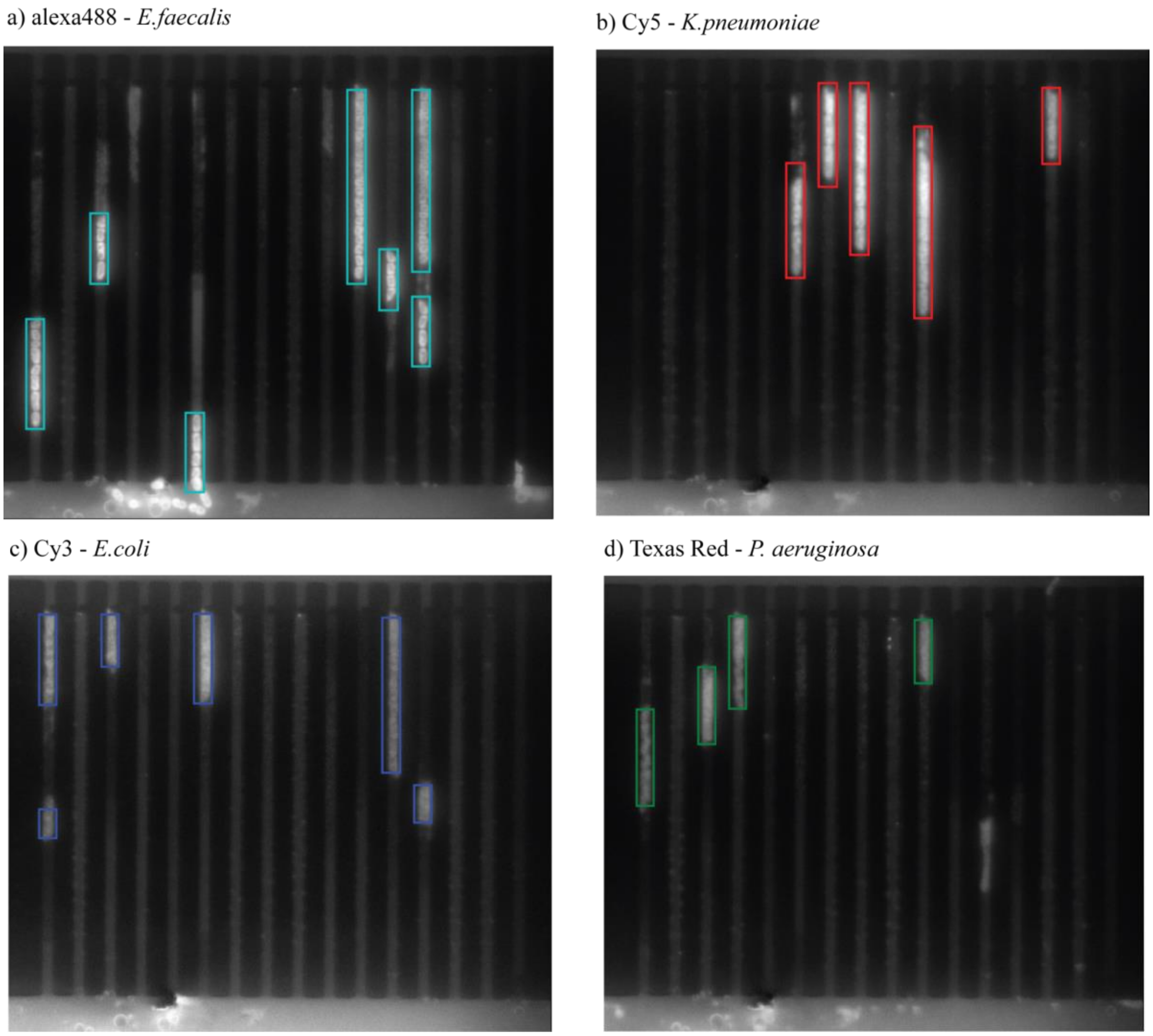


***Supplementary Figure 4:*** *Four fluorescent channel images corresponding to each FISH probe/species with bounding boxes around regions where cells are. a) Alexa 488 - E. faecalis b) Cy5 - K. pneumoniae c) Cy3 - E. coli d) Texas Red – P. aeruginosa.*

### Section 4: Cell tracking

Cell tracking involves linking cells from one frame to the next while they are growing in the mother-machine channels. A cell from frame at time *t* was compared against all the cells in time frame *t*+1, to generate a similarity score. This similarity score was obtained using a siamese network[5] whose inputs are “blob” images and blob properties like centroid, eccentricity, length, area, etc (**SI fig 5a**). The training data for this network was obtained by manually linking cells from one frame to the next using a GUI, as shown in **SI fig 5b**. A total of 212 frames were used in the training data, containing 3274 links. The training data was balanced to include cells that were linked and not-linked in equal proportions. Training data contained different species in equal proportions. Dividing cells were also present in the training data. 20 % of training data was set aside for validation.

The network loss function was set to minimize the difference between scores of similar pairs of objects (including mother-daughter splits) and to maximize the difference between scores of dissimilar pairs of objects. The exact loss function used is described in[6]. The margin in the loss function was set to 10. Cells that were similar were scored as close to 0 as possible and cells that were dissimilar were scored with values greater than 10. The training and validation loss curves are shown in **SI fig 5c**.

At test-time, a cell from frame at time *t* was compared to all the cells at time *t+1* to obtain similarity scores. **SI fig 5d** shows similarity scores between 2 consecutive frames. Low scores correspond to similar blobs. The lowest similarity scores for each blob is highlighted in blue in the score’s matrix. When a cell splits into two cells, their scores are very similar. The scores of a mother cell and its daughters are highlighted in red. An overall threshold (equal to 5.0) of similarity scores was set to exclude cells that were not similar to each other, and link cells that were similar. A second lower threshold (equal to 1.5) was set to allow for linking mother-daughter pairs. **SI fig 5d** shows links between two frames using such thresholds. The tracks that have rapid increase in area (over 40%) from one frame to the next were removed from the linking, when generating lineages. This helped in removing segmentation errors. Tracks were generated by stitching the series of links connecting blobs. **SI fig 5e**, shows different tracks on a stack of images from a single mother machine channel.

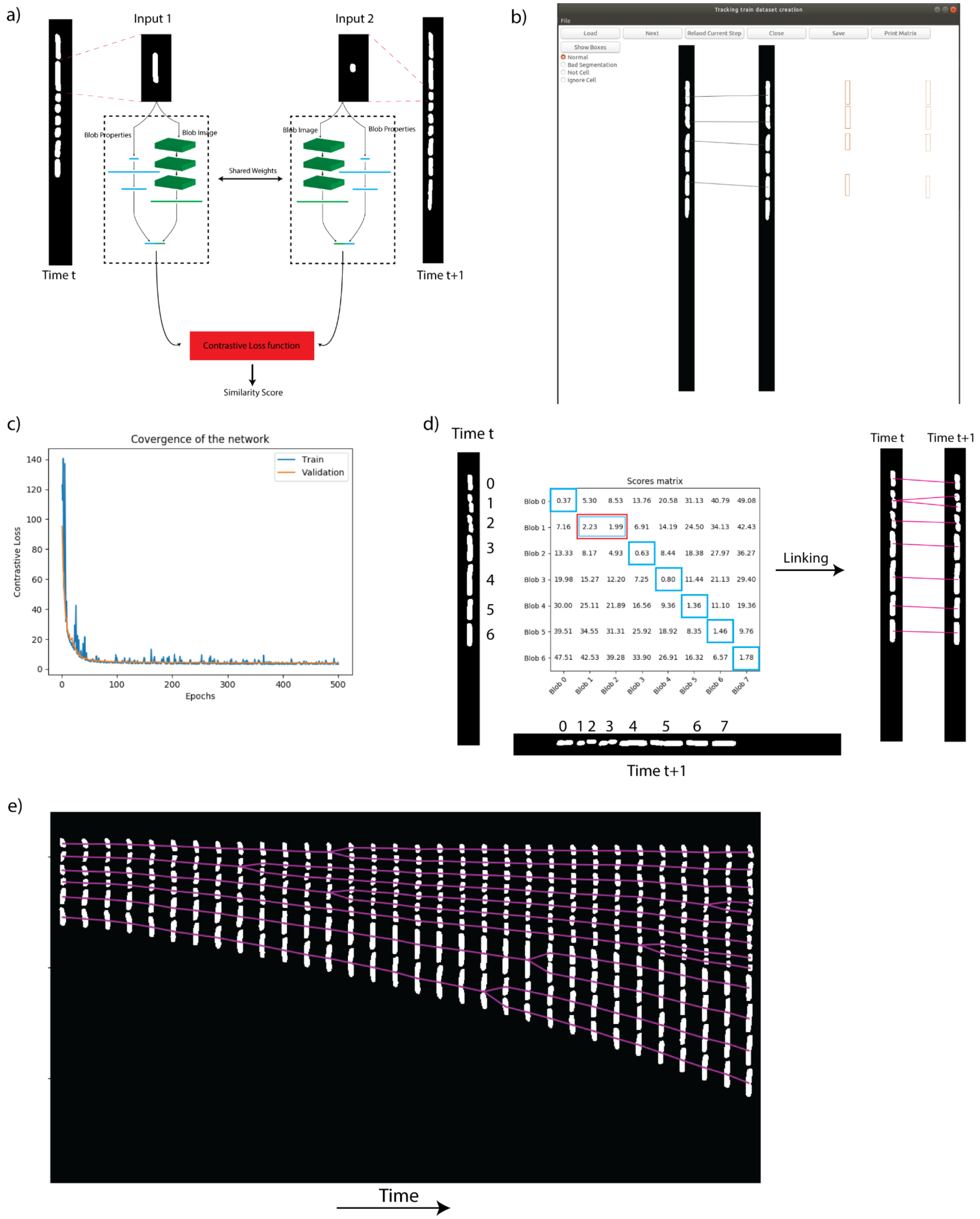


***Supplementary Figure 5:*** *a) Siamese network architecture b) GUI for generating training data. c) Training and validation loss curves d) Similarity scores between two frames and corresponding links e) Cell tracks from a single mother-machine channel.*

### Section 5: Species assignment for tracks using FISH data

The tracks ending in the last frame before fixing the cells were labeled with their species names obtained from the FISH data. This labeling was done based on which bounding boxes of the fluorescent data the centroids of the last blob in the track fell in. These species labels were rolled back in time to $t = 0$, thus labeling all the tracks since the beginning of the experiment.

**SI fig 6** shows a few of these labeled tracks in different colors for different species.

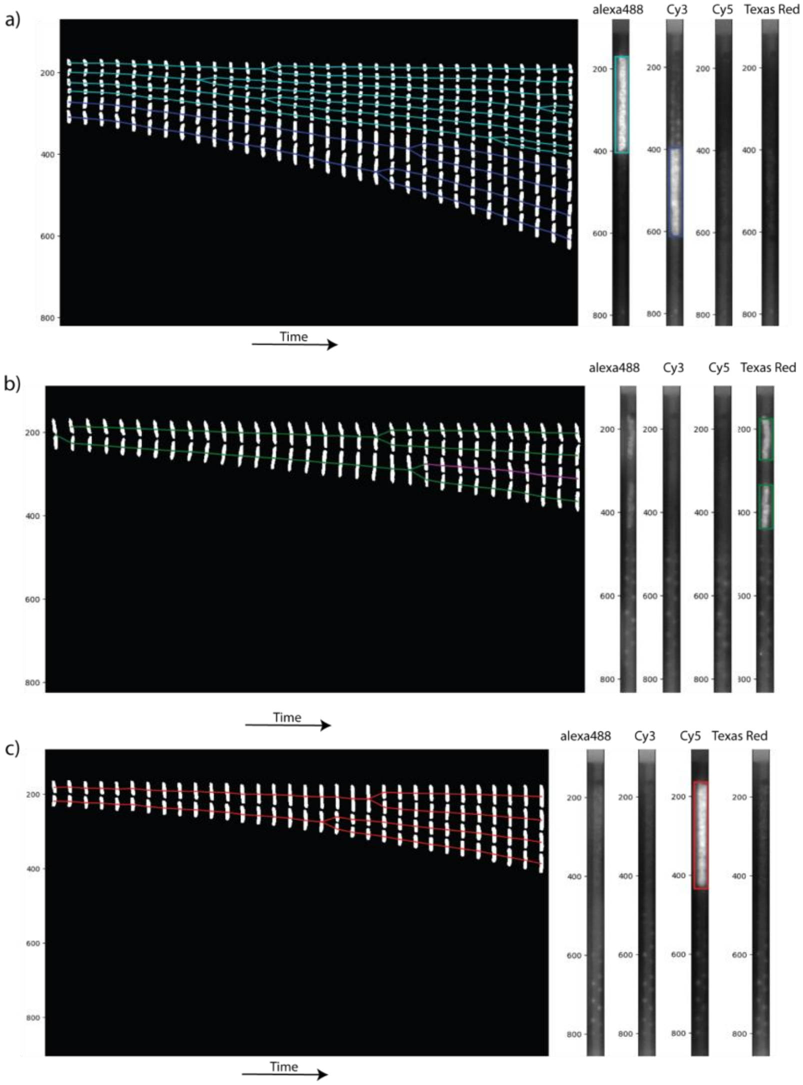


***Supplementary Figure 6:*** *Tracks labeled with different colors for different species a) E.faecalis (cyan) and E.coli(blue) in the same channel b) P.aeruginosa (green) and track with undetermined species(magenta) c) K.pneumoniae(red)*

### Section 6: Multi-Channel fluorescence Species assignment using PCA

**SI fig 7** shows images of single species loaded in four different chips and the images in four fluorescent channels after performing combinatorial FISH assay. It can be seen from the figure that the combination of two probes(colors) represent one single species.

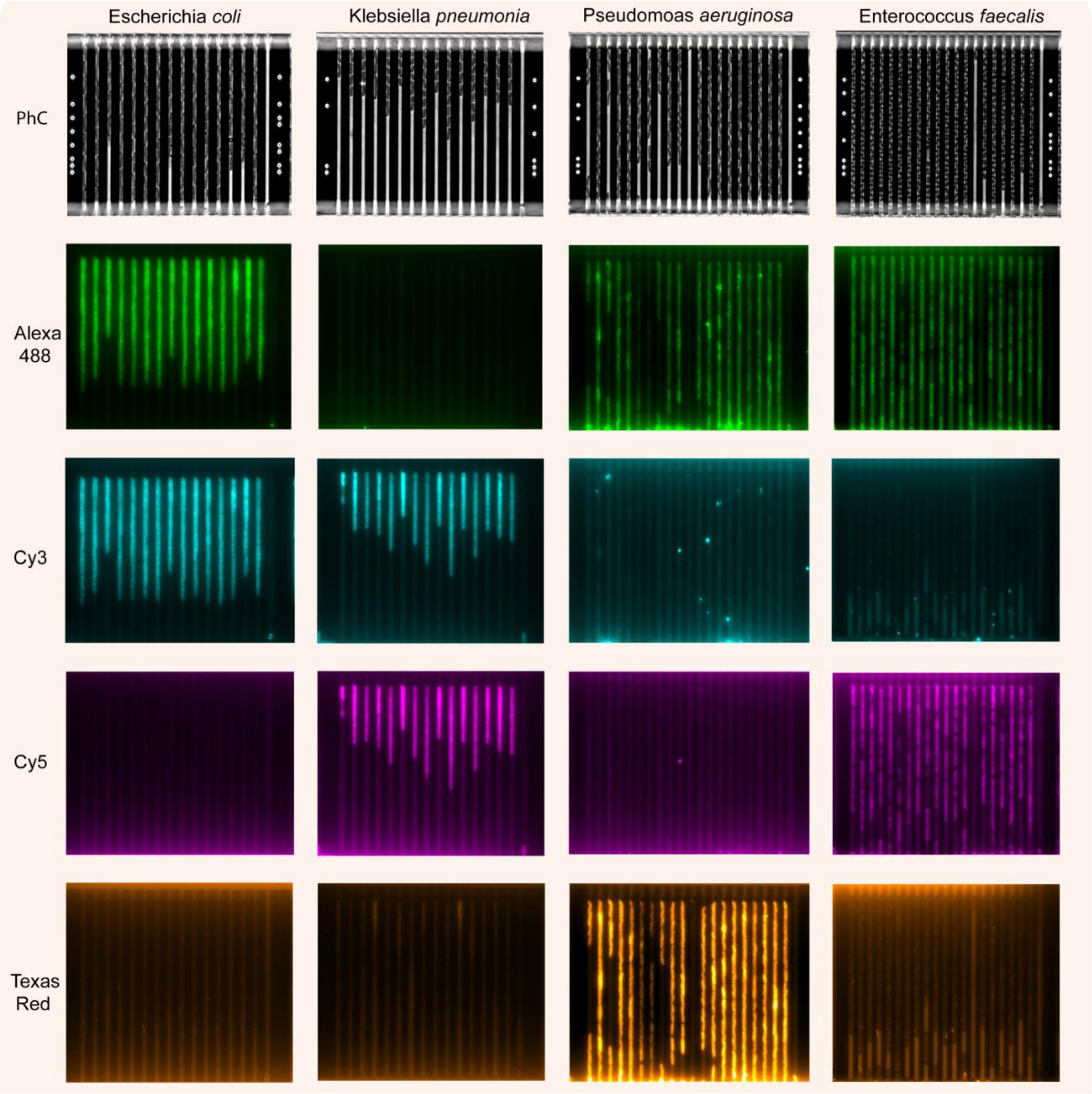


***Supplementary Figure 7:*** *Example images of individual bacterial species when performed combinatorial FISH. After the hybridization step, images were captured in different channels (PhC, Alexa 488, Cy3, Cy5, and Texas Red channels). Depending on the detection probe hybridized to adapter sequences of the individual bacterial species, cells were expected to be visualized in two different channels. For example*

*Escherichia coli (Alexa 488 and Cy3), Klebsiella pneumonia (Cy3 and Cy5) Pseudomonas aeruginosa (Alexa 488 and Texas redr) and Enterococcus faecalis (Alexa 488 and Cy5).*

In a typical experiment mixed species samples are loaded in the chip and combinatorial FISH assay is performed to get images such as the one included in the **main text (Fig 4b)**. As shown in **SI fig 7**, the adapter sequences and fluorescent probes are designed to bind to each species uniquely in two different fluorescent channels. **SI fig 8a,** shows the four fluorescence images of a single channel. This channel contains three species *P. aeruginosa, E.coli* and *K.pneumoniae* obtained using the combinations of fluorescence signals described in the **SI fig 7**. To assign species labels in individual mother machine channels, k-means clustering was performed on the average fluorescent signal along the width of the channel in all four fluorescence images. Data from 40 channels was used in this clustering. To visualize the clusters and assign each cluster with a species label, PCA was performed. Based on the combinations of signals observed in **SI fig 7**, each cluster's centroid was determined and labelled with species names. **SI fig 8b** shows different clusters of the first two principal components using data from 40 mother-machine channels. **SI fig 8c** shows the cluster to which the fluorescence data belongs to plotted against the length of the channel. From **SI fig 8a, b and c** it can be concluded that a unique assignment of species is achievable by resolving the data using PCA and the clustering using k-means. When presented with a new fluorescence signal data point one can determine the species based on distance to the nearest cluster's centroid.

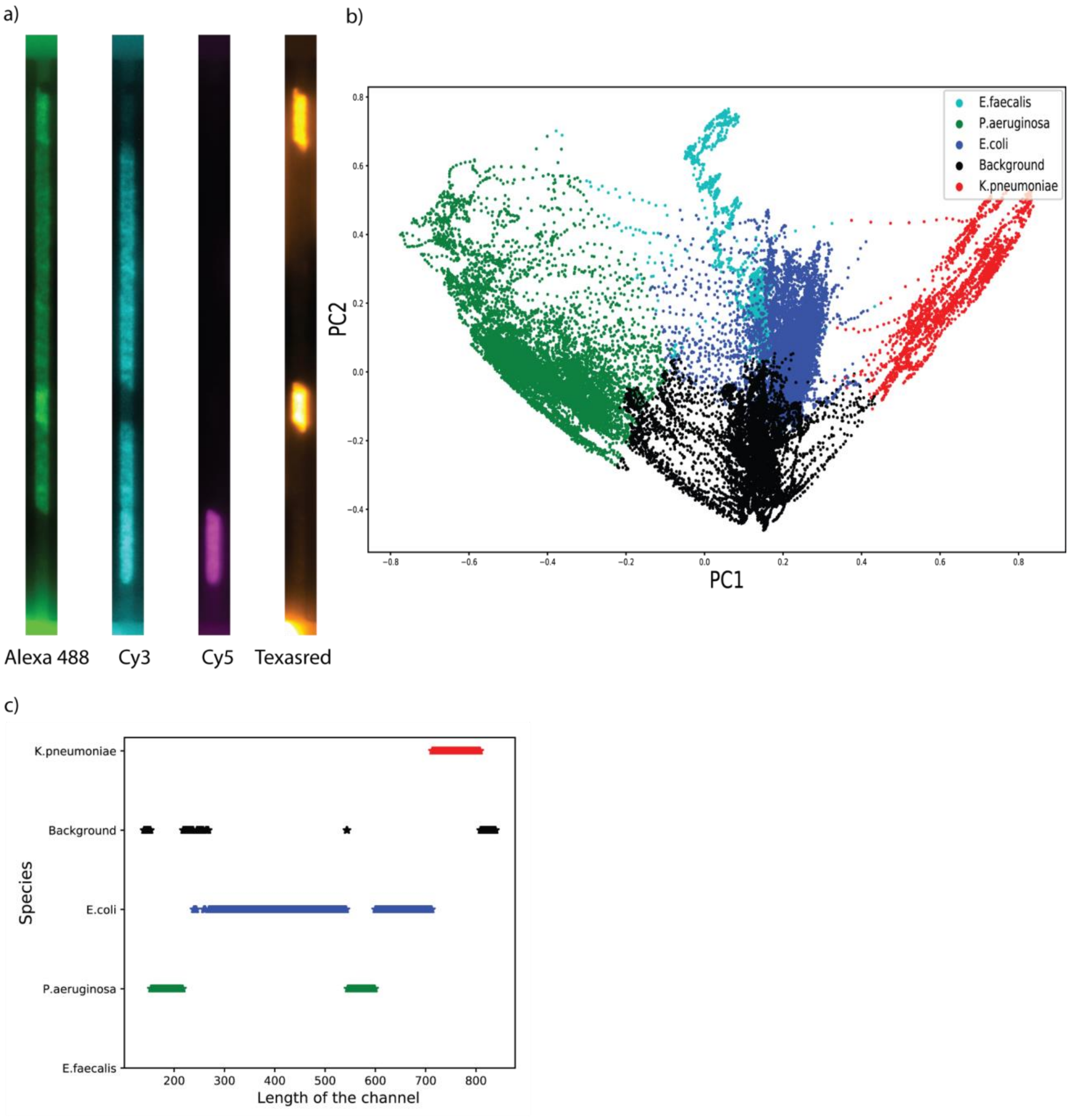


***Supplementary Figure 8:*** *PCA to assign species. a) Fluorescence Imaging in four channels of a single trap in the mother machine b) First two principal components of the four-dimensional mean fluorescence signal along the length of the channel clustered using K-means (n=5 clusters, 4 for species and 1 for background). c) Species-wise cluster numbers as we move along the length of the channel shown in (a).*

### Section 7: Growth Curve estimations

Growth rates were calculated by fitting exponential functions on the areas of cells in a single track on a rolling window of 5 frames, corresponding to 10 min. For each timepoint in the experiment, growth rates were collected from all the existing labeled tracks in that frame that existed for a 5 frame window. These growth rates were averaged to get the mean growth rate of that particular species. Growth rates were collected for both the reference and the antibiotic treated population. The normalized growth rate was calculated as the ratio of growth rates with respect to the treatment population. The plots in **SI fig 9a, 9b, 9c, 9d** show the normalized growth rates of all the four species used in an experiment along with their respective standard error of the mean. **SI fig 9e** shows the normalized growth rates of all 4 species plotted in the same graph for comparison. **SI fig 9f** shows the pooled normalized growth rates, i.e., growth rates without species identification. **SI fig 9g** shows the number of cells that went into tracking as a function of time and **SI fig 9h** shows the distribution of different species in the channels of the mother machine device.

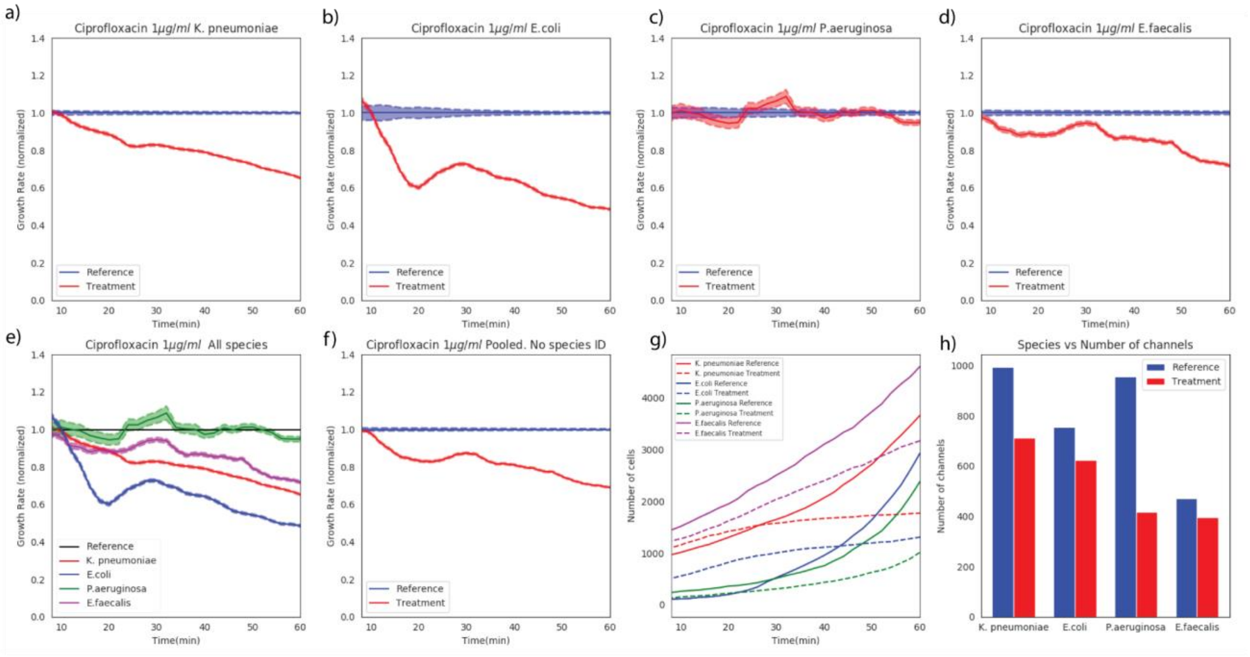


***Supplementary Figure 9:*** *A panel showing results of one experiment.*

### Section 8: Species wise probe sequence tables

**Table 1:** Oligonucleotides probes used for FISH

| **Target species** | **Sequence (5’ - 3’)** | **Fluorophore** | **Conjugation** | **Target site** | **Excitation** | **Emission** | **Vendor** |
|---|---|---|---|---|---|---|---|
| *Escherichia coli* | GCA TAA GCG TCG CTG CCG | Cy3/TYE 563 | 5’ | 23S *r*RNA | 530/11 | 575/19 | IDT |
| *Klebsiella pneumoniae* | CCT ACA CAC CAG CGT GCC | Cy5/TYE 665 | 5’ | 23S *r*RNA | 642/10 | 684/24 | IDT |
| *Pseudomonas aeruginosa* | TCT CGG CCT TGA AAC CCC | Texas Red | 5’ | 23S *r*RNA | 585/11 | 625/15 | IDT |
| *Enterococcus faecalis* | GAA AGC GCC TTT CAC TCT TAT GC | Alexa 488 | 5’ | 16S *r*RNA | 473/10 | 524/24 | IDT |

**Table 2:** Detection probes used for combinatorial FISH

| **Detection Probe** | **Sequence (5’ -3’)** | **Fluorophore** | **Conjugation** | **Excitation** | **Emission** | **Vendor** |
|---|---|---|---|---|---|---|
| DP 1 | TCATTGTCTCACTGCATTCG | Cy3/TYE 563 | 5’ | 530/11 | 575/19 | IDT |
| DP 2 | GTAAACAATGTGAAGCTCGG | Cy5/TYE 665 | 5’ | 642/10 | 684/24 | IDT |
| DP 3 | TGGTATGGTCACTTGGTATG | Texas Red | 5’ | 585/11 | 625/15 | IDT |
| DP 4 | ACTGTCGCTCGCTCAATCTG | Alexa 488 | 5’ | 473/10 | 524/24 | IDT |

**Table 3:** oligonucleotide sequences used for the combinatorial FISH

| **Name** | **Oligo Sequence 5’-3’ (Barcode-target-Barcode)** | **Target species** | **Vendor** |
|---|---|---|---|
| B1*Esc*B4 | CGAATGCAGTGAGACAATGAGCATAAGCGTCGCTG CCGCAGATTGAGCGAGCGACAGT | *Escherichia coli* | IDT |
| B1*Kpn*B2 | CGAATGCAGTGAGACAATGACCTACACACCAGCGT GCCCCGAGCTTCACATTGTTTAC | *Klebsiella pneumoniae* | IDT |
| B4*Pse*B3 | CAGATTGAGCGAGCGACAGTTCTCGGCCTTGAAAC CCCCATACCAAGTGACCATACCA | *Pseudomonas aeruginosa* | IDT |
| B2*Enf*B4 | CCGAGCTTCACATTGTTTACGAAAGCGCCTTTCACT CTTATGCCAGATTGAGCGAGCGACAGT | *Enterococcus faecalis* | IDT |

**Table 4:** Barcode sequences used to link with the target sequence

| Barcode sequence name | Sequence (5'-3') | Vendor |
|---|---|---|
| BS 1 | CGAATGCAGTGAGACAATGA | IDT |
| BS 2 | CCGAGCTTCACATTGTTTAC | IDT |
| BS 3 | CATACCAAGTGACCATACCA | IDT |
| BS 4 | CAGATTGAGCGAGCGACAGT | IDT |

#### Section 9: Supplementary Movies

Movie S1-S5: An example of a time-lapse movie and corresponding FISH image of mixed species growing in the microfluidic chip supplied with MH media without the antibiotics (S1) and with antibiotics Van (S2), CIP (S3), Gen (S4) and NIT (S5). This movie is generated from the time-lapse phase contrast images with time intervals of 120 sec and playback speed of the video is 5 fps.
