## Supplementary movies for "Rapid antibiotic susceptibility testing and species identification for mixed infections"

### Slide 1
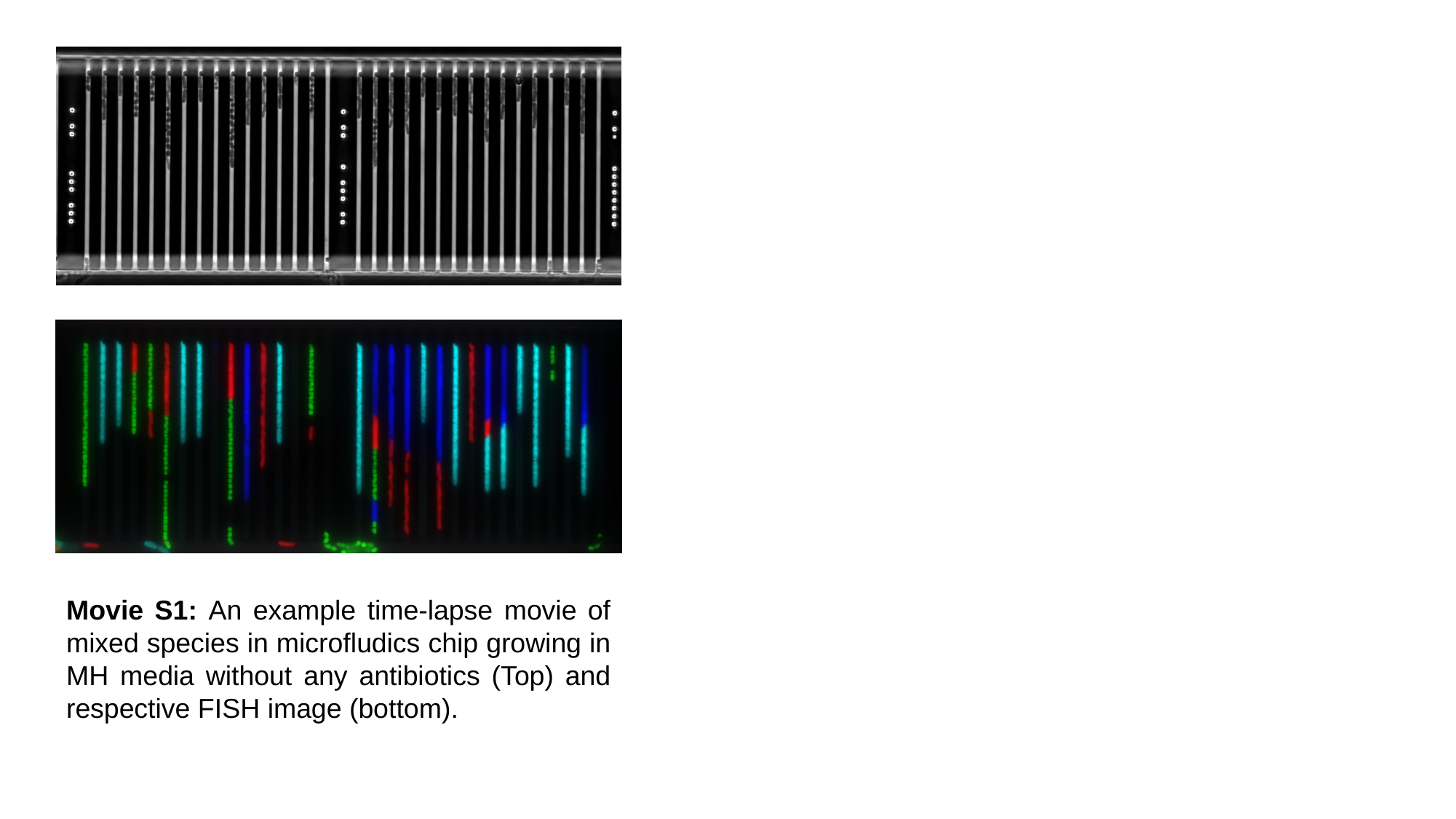

Movie S1: An example time-lapse movie of mixed species in microfludics chip growing in MH media without any antibiotics (Top) and respective FISH image (bottom).

### Slide 2
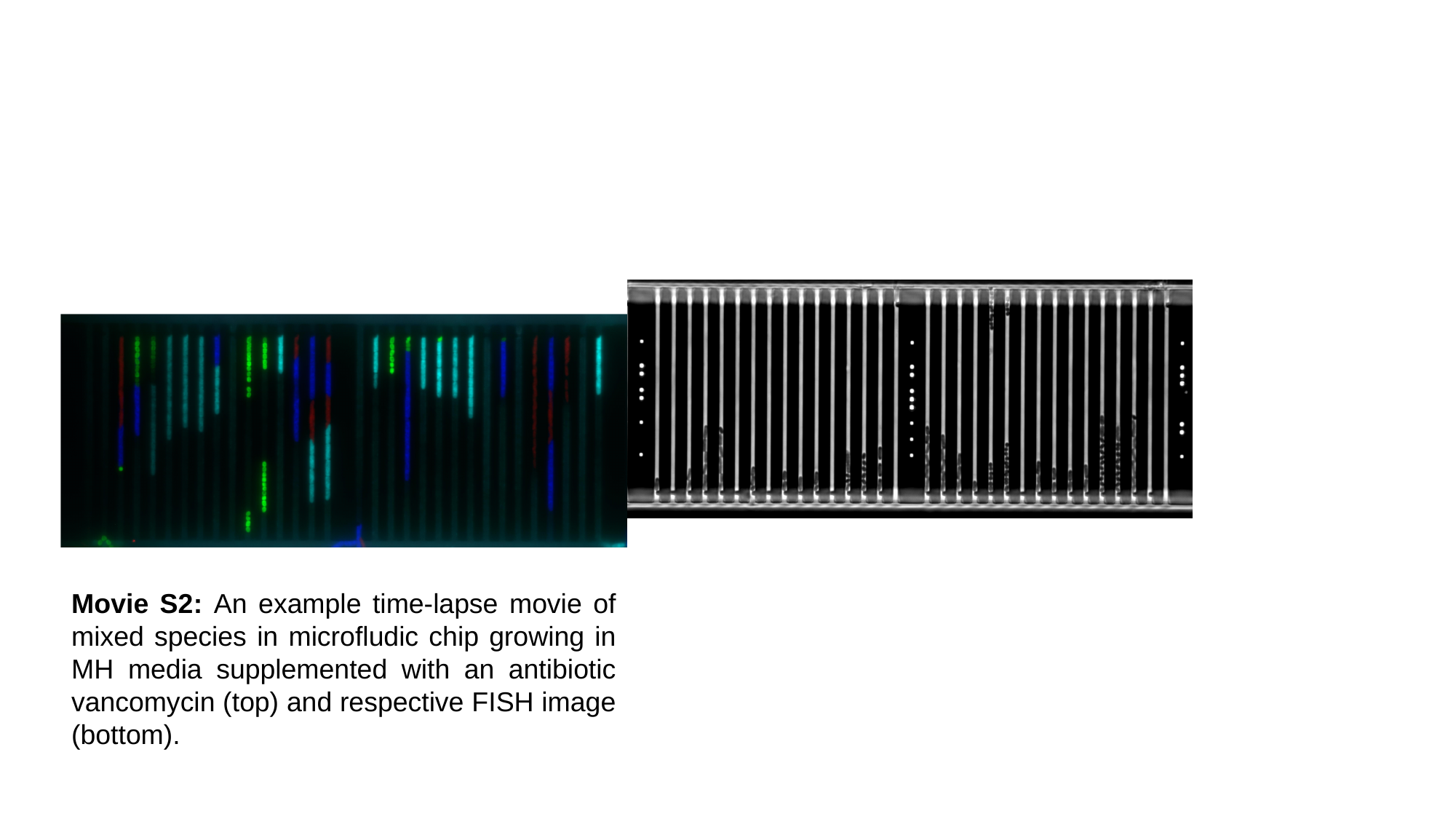

Movie S2: An example time-lapse movie of mixed species in microfludic chip growing in MH media supplemented with an antibiotic vancomycin (top) and respective FISH image (bottom).

### Slide 3
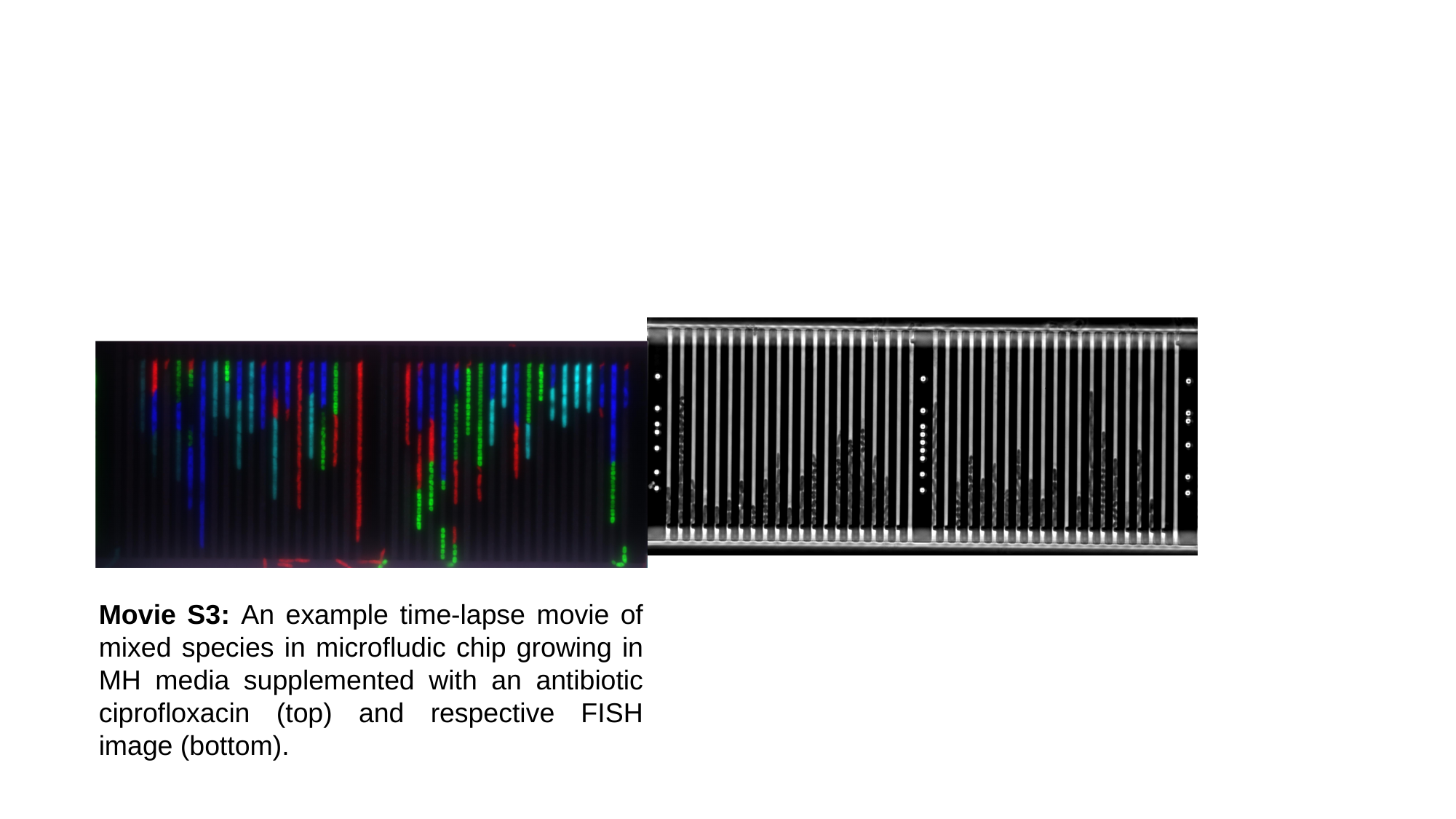

Movie S3: An example time-lapse movie of mixed species in microfludic chip growing in MH media supplemented with an antibiotic ciprofloxacin (top) and respective FISH image (bottom).

### Slide 4
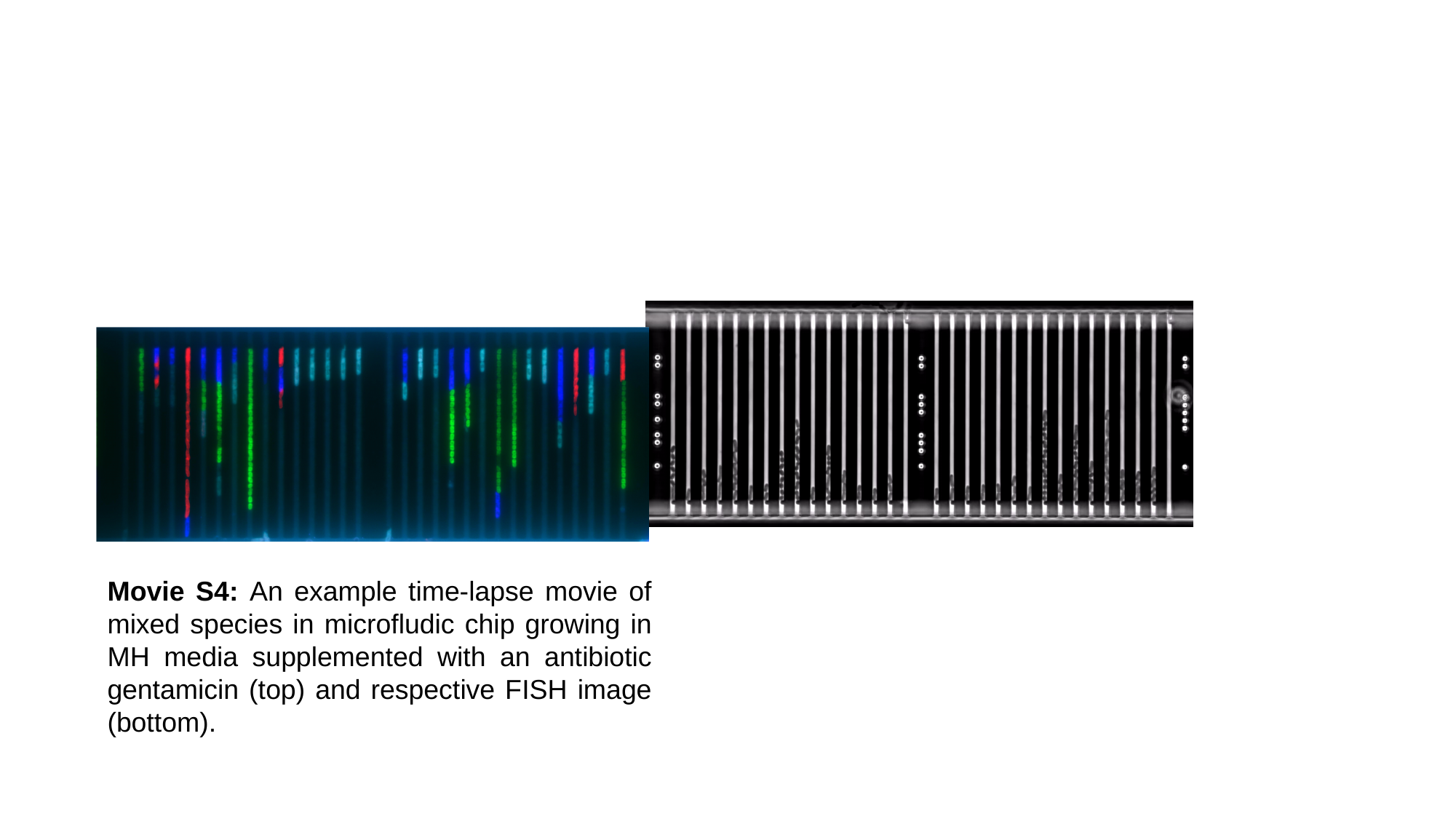

Movie S4: An example time-lapse movie of mixed species in microfludic chip growing in MH media supplemented with an antibiotic gentamicin (top) and respective FISH image (bottom).

### Slide 5
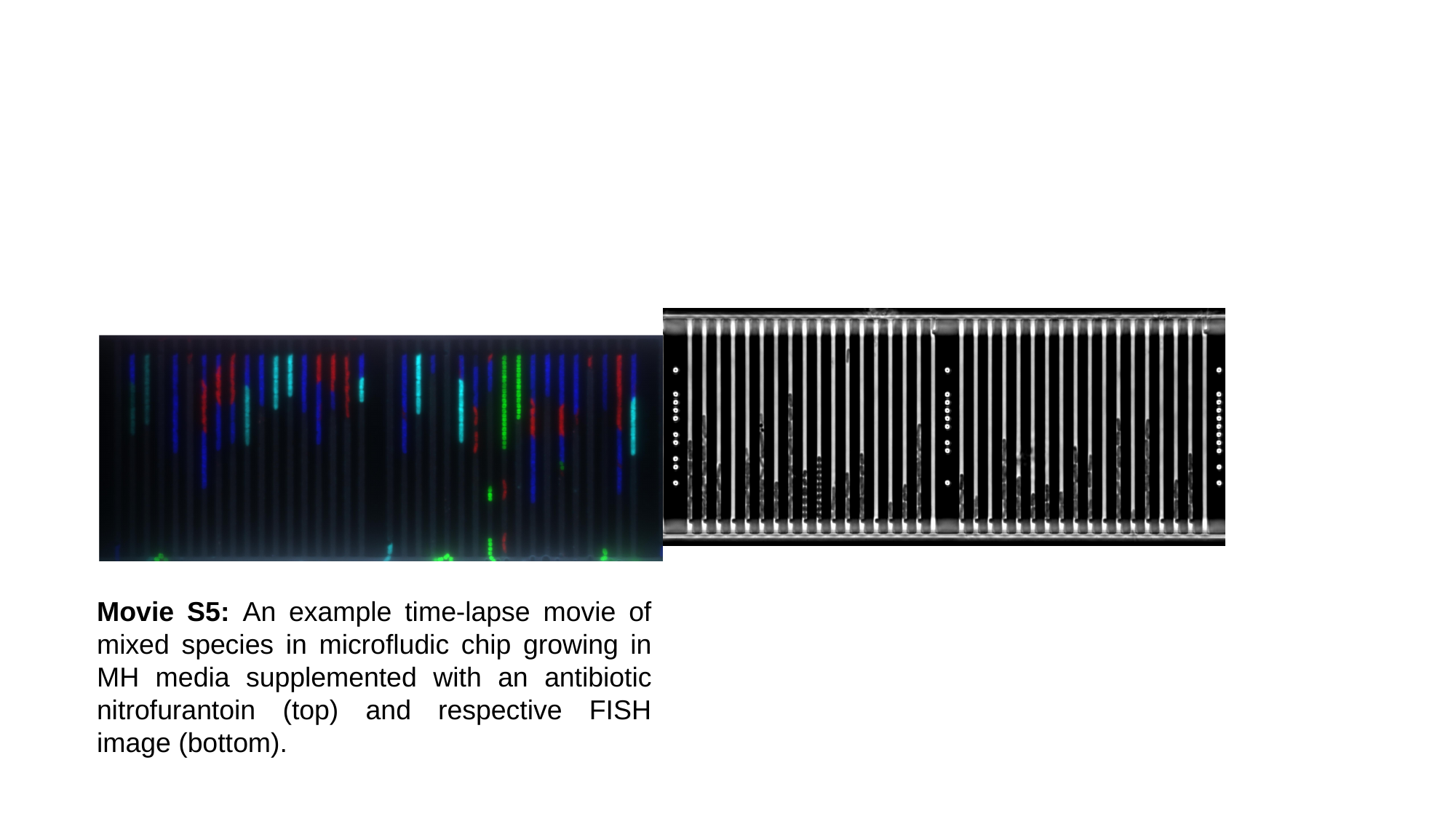

Movie S5: An example time-lapse movie of mixed species in microfludic chip growing in MH media supplemented with an antibiotic nitrofurantoin (top) and respective FISH image (bottom).
